# DISTINCT ROLES OF VIRAL US3 AND UL13 PROTEIN KINASES IN HERPES VIRUS SIMPLEX TYPE 1 (HSV-1) NUCLEAR EGRESS

**DOI:** 10.1101/2023.03.20.533584

**Authors:** Masoudeh Masoud Bahnamiri, Richard J. Roller

**Author notes:** Corresponding author 3-432 Bowen Science Building 51 Newton Road Iowa City, IA 52242.

## Abstract

Herpesviruses transport nucleocapsids from the nucleus to the cytoplasm by capsid envelopment into the inner nuclear membrane and de-envelopment from the outer nuclear membrane, a process that is coordinated by nuclear egress complex (NEC) proteins, pUL34, and pUL31. Both pUL31 and pUL34 are phosphorylated by the virus-encoded protein kinase, pUS3, and phosphorylation of pUL31 regulates NEC localization at the nuclear rim. pUS3 also controls apoptosis and many other viral and cellular functions in addition to nuclear egress, and the regulation of these various activities in infected cells is not well understood. It has been previously proposed that pUS3 activity is selectively regulated by another viral protein kinase, pUL13 such that its activity in nuclear egress is pUL13-dependent, but apoptosis regulation is not, suggesting that pUL13 might regulate pUS3 activity on specific substrates. We compared HSV-1 UL13 kinase-dead and US3 kinase-dead mutant infections and found that pUL13 kinase activity does not regulate the substrate choice of pUS3 in any defined classes of pUS3 substrates and that pUL13 kinase activity is not important for promoting de-envelopment during nuclear egress. We also find that mutation of all pUL13 phosphorylation motifs in pUS3, individually or in aggregate, does not affect the localization of the NEC, suggesting that pUL13 regulates NEC localization independent of pUS3. Finally, we show that pUL13 co-localizes with pUL31 inside the nucleus in large aggregates, further suggesting a direct effect of pUL13 on the NEC and suggesting a novel mechanism for both UL31 and UL13 in the DNA damage response pathway.

IMPORTANCE

Herpes simplex virus infections are regulated by two virus-encoded protein kinases, pUS3 and pUL13, which each regulate multiple processes in the infected cell, including capsid transport from the nucleus to the cytoplasm. Regulation of the activity of these kinases on their various substrates is poorly understood, but importantly, kinases are attractive targets for the generation of inhibitors. It has been previously suggested that pUS3 activity on specific substrates is differentially regulated by pUL13 and, specifically, that pUL13 regulates capsid egress from the nucleus by phosphorylation of pUS3. In this study, we determined that pUL13 and pUS3 have different effects on nuclear egress and that pUL13 may interact directly with the nuclear egress apparatus with implications both for virus assembly and egress and, possibly, the host cell DNA-damage response.

## INTRODUCTION

Herpesviruses use a sequential capsid envelopment and de-envelopment pathway for the transport of the capsid across the nuclear envelope (1, 2). This nuclear egress pathway is mediated by a heterodimer of two viral proteins, pUL31/pUL34, called the nuclear egress complex (NEC), which is conserved among all herpesviruses (3, 4, 5). In infected cells and in cells that ectopically express only pUL31 and pUL34, the two proteins self-assemble into heterodimers that localize at the nuclear rim (3, 6, 7, 8, 9, 10). Expression of pseudorabies virus (PRV) pUL31 and pU34 in the absence of other viral proteins results in the nuclear membrane vesiculation into the perinuclear space (PNS) (11), and purified, recombinant HSV-1 and PRV pUL31 and pUL34 can induce budding into giant unilamellar vesicles (GUVs) or liposomes (12, 13). Purified NECs of HSV and PRV crystallize as arrays of hexamers of the NEC heterodimer, and membrane budding *in vitro* by the HSV or PRV NEC is associated with the formation of hexameric arrays, indicating that the NEC can spontaneously self-associate into these higher-order complexes. Furthermore, mutations in the HSV-1 NEC that are predicted to disrupt hexamer-forming interactions also inhibit budding, suggesting that the formation of higher-order NEC complexes drives the membrane curvature (14). Clearly, regulation of NEC self-association is critical for the regulation of membrane curvature. Interestingly in wild-type HSV-1-infected cells, budding of empty vesicles into the perinuclear space (PNS) is rarely observed, suggesting the NEC self-association that leads to budding is negatively regulated and triggered by the association of capsids with the INM.

One regulator of NEC self-association is the virus-encoded protein kinase, pUS3, which is conserved within the alphaherpesvirus subfamily, including HSV and PRV. When pUS3 is absent or catalytically inactive, the localization of the NEC changes from a roughly evenly distribution on the nuclear envelope to the accumulation of punctate structures, suggesting that pUS3 negatively regulates the self-association of the NEC. These same pUS3 mutants also cause the accumulation of capsids in the PNS, implicating pUS3 in the de-envelopment of capsids from the PNS (15, 16, 17, 18). Interestingly, N-terminal phosphorylation of pUL31 by pUS3 is thought to be one critical event in promoting de-envelopment, as serine mutations to alanine in this region of pUL31 also result in capsid accumulation in the PNS (19), suggesting the possibility that pUS3 phosphorylation of UL31 drives the de-oligomerization of the NEC and release of the capsid into the cytoplasm. Interestingly, a recent study reported that pUL21 recruits the protein phosphatase 1 (PP1) and mediates the dephosphorylation of UL31, which might oppose the effects of US3 kinase activity in the regulation of NEC (20). It is likely that

UL21-mediated dephosphorylation of pUL31 facilitates NEC self-association that leads to membrane budding into the nuclear membrane.

Nuclear shape and nuclear envelope rigidity are maintained by the nuclear lamina, a network of intermediate filament family proteins that underlies the INM and is connected to it by interaction with integral membrane lamin-associated proteins (LAPs) (21, 22, 23, 24). Access of capsids to the INM for envelopment and deformation of the nuclear envelope for budding are both thought to be inhibited by the intact nuclear lamina (2, 25). In herpesvirus-infected cells, the NEC recruits protein kinases to phosphorylate both lamins and LAPs, resulting in changes to nuclear shape and distribution of the lamina proteins in the nuclear membrane (15, 26, 27, 28, 29, 30, 31, 32). pUS3 is one of these kinases, and it both promotes and regulates nuclear lamina redistribution. pUS3 phosphorylates both lamin subunits and (LAPs), promoting their redistribution, but its ability to inhibit NEC self-association (33) also regulates lamina dissociation such that when pUS3 is absent or catalytically inactive, large gaps form in the nuclear lamina at the sites of NEC self-association (7, 15).

In addition to the regulation of nuclear egress, pUS3 is a potent virulence factor involved in the regulation of many different cellular processes during HSV-1 infection. pUS3 upregulates at least 23 known events in the infected cell (34), including inhibition of the apoptosis (35, 36, 37, 38, 39, 40, 41, 42, 43), rearrangement of the microtubule network to control infected cellmorphology (43, 44), evasion of host-cell immune response in various ways (45, 46, 47), enhancement of viral gene expression (48), stimulation of protein translation (49), promotion of viral spread, and regulation of intracellular trafficking of viral and cellular factors (50, 51, 52). Deletion of US3 in mouse models of HSV infection results in severe attenuation (53). In cell culture, however, US3 deletion results in virus replication defects in a cell type-dependent manner. While previous reports showed at most a ten-fold single-step growth defect associated with US3-null or US3-catalytically inactive mutants, recent studies from our laboratory demonstrated that US3 kinase-inactive mutants show a roughly 250-fold decrease in virus production in physiologically more relevant cell lines (33). Epistasis mutagenesis analysis from the same study also suggested that about a ten-fold growth defect arises as a result of pUS3 failure to do nuclear egress functions.

The multiple and diverse functions attributed to pUS3 suggest that its activity in these processes might be differentially regulated, and indeed, Kato et al. have presented results suggesting that pUS3 activity, specifically in nuclear egress, might be regulated by another viral protein kinase, pUL13 (54). pUL13 is a conserved herpesviruses protein kinase (CHPK), whose homologs in all herpesviruses have some overlap in function with cyclin-dependent protein kinases (55, 56). pUL13 homologs in beta and gammaherpesvirus regulate nuclear egress in a different but overlapping way compared to pUS3 in alpha-herpesviruses. Inhibition of CHPKs expression results in a significant viral growth defect in human beta and gammaherpesviruses (55, 57, 58, 59, 60). This growth defect is associated with a reduction in the levels of NEC proteins that results in impaired nuclear egress (55, 57, 61, 62). However, in alphaherpesviruses, pUL13 is not essential and has cell-type-specific effects on viral replication in cell culture (63, 64, 65, 66, 67, 68, 69). It is possible that the significance of UL13 in α- herpesviruses replication and nuclear egress is diminished because some characteristic CHPK activities have been assumed by pUS3. pUL13 phosphorylates pUS3 both in vivo and in vitro, and while pUL13 regulates NEC distribution in a manner similar to US3, it does not regulate apoptosis in a manner similar to US3 (54), suggesting that pUL13 might regulate NEC localization by phosphorylation of pUS3 without affecting some other activities of pUS3. Here, we further explored the pUL13/pUS3 relationship in several ways. First, we characterized the effect of pUL13 kinase activity on NEC organization. We found that the NEC self-association formed in pUL13 mutant infection is quite different from pUS3 mutants and that pUL13 mutants show no obvious de-envelopment defect. Second, we tested the hypothesis that pUL13 regulates the substrate specificity of US3 by testing representatives of different groups of pUS3 substrates, and we found no pUS3 substrate whose phosphorylation was dependent on pUL13 catalytic activity. Finally, while the results of Kato et al. showed an interesting correlation between failure to phosphorylate pUS3 and the formation of NEC self-association, it did not directly show that these events were causally related. We observed that mutation of pUL13 phosphorylation motifs on pUS3, individually or in aggregate, had no effect on NEC localization, suggesting that pUL13 might have a direct effect on the NEC. In support of this, we further found that pUL13 directly interacts with pUL31.

## MATERIALS & METHODS

### Cell lines

Vero cells were maintained in Dulbecco’s modified Eagle medium (DMEM) (Thermo Fisher Scientific) supplemented with 5% bovine calf serum (BCS) and penicillin-streptomycin (P/S). HaCaT (kind gift from David Johnson) and 293T cells were maintained in DMEM supplemented with 10% fetal bovine serum (FBS) and P/S. SH-SY5Y (kind gift from Stevens Lewis at the University of Iowa) was maintained in Opti-MEM supplemented with 5% FBS, sodium pyruvate, non-essential amino acid, and P/S. All the cells were cultured in a humidified incubator at 37°C with 5% CO_2_.

### Construction of recombinant viruses

The HSV-1(F) bacterial artificial chromosome (BAC) described previously by (70) (kind gift from Yasushi Kawaguchi at the University of Tokyo) was used for the construction of all the recombinant viruses (70). The recombinant virus with a point mutation at residue 220 in US3 was constructed using gentamycin cassette insertion and excision as previously described (71). For each recombinant virus, the mutagenesis insertion DNA fragment that was designed as two fragments and amplified by PCR are as described in Table 2. All the recombinant viruses were rescued by transfecting BAC DNA into Vero cells using lipofectamine DNA transfection reagent (Millipore-Sigma) according to the manufacturer’s instructions. The HSV-1(F) BAC-derived UL13_K176A_ was generated by creating a point mutation at invariant lysine residue 176 in UL13. The HSV-1(F) BAC-derived US3_S47A_, US3_S139A_, US3_S336A_, and US3_S413A_ mutant recombinant viruses were generated by creating a point mutation at residue 47, 139, 336, and 413 in US3 respectively. US3_S139A_ BAC was used as a backbone to introduce 3 more amino acid substitutions sequentially to build up US3SA4. Genomes were isolated from all recombinant viruses and Illumina sequencing was performed by SeqCenter LLC as previously described (72). Illumina reads were mapped onto the pYeBAC102 genome as previously described (72). The only differences observed were the intended, engineered mutations and heterogeneity in the length of G/C homopolymer runs in intergenic sequences.

### Plasmids

A pcDNA3 plasmid carrying N-terminally HA-tagged UL34 (pRR1385) and a pcDNA3 plasmid carrying C-terminally FLAG-tagged UL31 were described previously (71).

A plasmid expressing C-terminally EGFP-tagged UL13 (pRR1404) was constructed by the amplification of the UL13 coding sequence with primers (FW 5’- GAGCTCAAGCTTCGAATTCTATGGATGAGTCCCGCAGACAGC—3’ and RV, 5’ GTATGGCTGATTATGATCAGTCACGACAGCGCGTGCC—3’) and insertion into the pEGFP- C1 vector by Gibson assembly.

### Immunofluorescence assay

For assay of infected cells, Vero, HaCaT, or SH-SY5Y cells grown on coverslips were infected with viruses at an MOI of 5 and fixed at 16 hpi. with 4% formaldehyde for 20 min followed by three washes with PBS. Transfected cells grown on coverslips were fixed 48 hours after transfection. Fixed cells were permeabilized and blocked with a 30-minute incubation in Immunofluorescence (IF) buffer (1X PBS, 0.5% Triton X-100, 0.5% sodium deoxycholate, 1% egg albumin, 0.01% NaN_3_) and then incubated for 1 h in primary antibody diluted in IF buffer. Coverslips were then washed twice with PBS and incubated for 1 h in fluorescently labeled secondary antibodies diluted 1:1000 in IF buffer. Primary antibodies for assay of infected cells were chicken polyclonal anti-UL34 (1:250) (71) and mouse monoclonal anti-LMNA/C (Santa Cruz Biotechnology), both diluted 1:500. Primary antibodies for assay of transfected cells were goat polyclonal anti-HA (1:500, GenScript), and mouse monoclonal anti-FLAG (1:500, Sigma-Aldrich). Confocal microscopy was performed on a Leica DFC7000T confocal microscope using Leica software. FIJI ImageJ software was used to quantify localization signals. The images shown are representative of experiments performed at least three times.

### Transfection

Vero cells grown on glass coverslips in 24 well culture plates were transfected with EGFP-tagged UL13, HA-tagged UL34, and FLAG-tagged UL31 plasmid constructs using of X-tremeGENE HP DNA transfection reagent (Millipore-Sigma) according to the manufacturer’s instructions. Transfected cells were fixed after 48 h.

### Transmission electron microscopy (TEM)

Confluent monolayers of Vero or HaCaT cells were infected with HSV-1(F) BAC, US3_K220A_, and two isolates of UL13_K176A_ or US3_S47A_ virus strains at an MOI of 5 and fixed 20 hpi by incubation in 2.5% glutaraldehyde in 0.1 M cacodylate buffer (pH 7.4) for 2 h. Following fixation in 1% osmium tetroxide, cells were washed in cacodylate buffer, and embedded in Spurr’s resin. Cells were cut into 95-nm sections that were mounted on grids, and stained with uranyl acetate and lead citrate to examine with a JEOL 1250 transmission electron microscope.

### Single-step growth measurement (SSG)

Replication of the HSV-1(F) BAC, US3_K220A_, UL13_K176A_, US3_S47A_, US3_S139A_, US3_S336A_, US3_S413A_, and US3_SA4_ virus strains on Vero, HaCaT, or SH-SY5Y cells were measured as previously described (4). Log-transformed data were used for statistical analysis

### Emerin hyperphosphorylation assay

HaCaT cells in 100 mm culture plates were either mock-infected or infected with HSV-1(BAC), US3_K220_, or UL13_K176A_, virus strains at an MOI of 5 for 16. Infected cells were washed and collected in phosphate-buffered saline (PBS) and nuclear envelope proteins were isolated as previously described (26). Cell lysates and nuclear envelope fractions were separated with 10% SDS-PAGE gels, blotted onto nitrocellulose membranes, and probed for emerin using mouse monoclonal anti-emerin (1:500) (Santa Cruz Biotechnology). The membranes were then incubated with the respective alkaline phosphatase-conjugated secondary antibodies for 1 hour and developed in western blotting (WB) development solution as previously described.

### Western blotting

HaCaT cells in 6-well culture plates were either mock-infected or infected with HSV-1(BAC), US3_K220_, or UL13_K176A_ virus strains, respectively, at an MOI of 5. At 18 hpi, infected cells were collected in phosphate-buffered saline (PBS) and lysed with RIPA buffer (50 mM Tris, pH 7.5, 150 mM NaCl, 1 mM EDTA, 1% Triton X-100). Cell lysates were separated with 10% SDS-PAGE gels, blotted onto nitrocellulose membranes, and probed for PRKAR2A using (mouse anti) rabbit anti-phospho-PKA substrates (1:500) (Cell Signaling), rabbit anti-phospho-Akt substrates (1:500) (Cell Signaling), mouse monoclonal anti-actin (Sigma-Aldrich), chicken polyclonal anti-UL34 (1:250) (4). The membranes were then incubated with the respective alkaline phosphatase-conjugated secondary antibodies for 1 hour and developed in western blotting (WB) development solution as previously described.

### US3 mobility shift assay

Vero or HaCaT cells in 6-well culture plates were either mock-infected or infected with HSV-1(BAC), US3-null (16), US3_K220_, UL13_K176A_, US3_S47A_, US3_S139A_, US3_S336A_, US3_S413A_ and US3_SA4_ virus strains, respectively, at an MOI of 5. At 18 hpi, infected cells were collected in phosphate-buffered saline (PBS) and lysed with RIPA buffer (50 mM Tris, pH 7.5, 150 mM NaCl, 1 mM EDTA, 1% Triton X-100). Nuclei were pelleted and discarded prior to solubilization in SDS sample buffer. Cell lysates were separated with 10% SDS-PAGE gels, blotted onto nitrocellulose membranes, and probed as previously described (4) using chicken polyclonal anti-UL34 (1:250)(4), rabbit polyclonal anti-HSV-1 US3 (kind gift from B. Roizman), and mouse monoclonal anti-actin (Sigma-Aldrich).

### Co-immunoprecipitation assay of gB

Vero cells in 100 mm culture plates were either mock-infected or infected with HSV-1(BAC), US3_K220_, or UL13_K176A_ virus strains at an MOI of 5. At 18 hpi, infected cells were collected in phosphate-buffered saline (PBS) and lysed with RIPA buffer (50 mM Tris, pH 7.5, 150 mM NaCl, 1 mM EDTA, 1% Triton X-100). The lysate was incubated with rabbit anti-gB diluted 1:1000 (kind gift from G. Cohen and R. Eisenberg) overnight at 4°C. Cells were precipitated using Dynabeads^TM^ protein A (Invitrogen) according to the manufacturer’s instructions. Cell lysates were separated with 10% SDS-PAGE gels, blotted onto nitrocellulose membranes, and probed for gB with rabbit anti-gB diluted 1:1000 (kind gift from G. Cohen and R. Eisenberg) or for phospo-gB with rabbit anti-phospho-PKA substrates diluted 1:500 (Cell Signaling).

### Co-immunoprecipitation assay of EGFP-UL13

293T cells in 100 mm culture plates were transfected with plasmids EGFP-tagged UL13 and FLAG-tagged UL31 plasmid constructs by use of PEI (polyethyleneimine) DNA transfection reagent (1:9) as previously described (71) (Roller et al. 2010). Cells were collected 48 hours post-transfection in phosphate-buffered saline (PBS) and lysed with RIPA buffer (50 mM Tris, pH 7.5, 150 mM NaCl, 1 mM EDTA, 1% Triton X-100). The lysates were sonicated, and the supernatants were incubated with 10 µl FLAG M2 Magnetic Beads (Milipore-Sigma) overnight at 4°C, washed with PI-RIPA, and eluted with FLAG peptide in PI-RIPA according to the manufacturer’s instructions (Milipore-Sigma). Cell lysates were separated with 10% SDS-PAGE gels, blotted onto nitrocellulose membranes, and probed for EGFP with rabbit anti-EGFP diluted 1:5000 (kind gift from C. Ellermier) or for FLAG with mouse anti-FLAG M2 diluted 1:500 (Milipore-Sigma).

### Quantification of the frequency of NEC aggregation

Randomly selected fields were used to take images of more than 100 cells’ nuclei via a confocal microscope. The frequency of the nucleus with aggregated NEC or disrupted LaminA/C was counted using FIJI ImageJ software.

### Nuclear morphology analysis

Nuclear contour ratios were calculated using cross-sectional images of nuclei of Vero and HaCaT uninfected cells or cells infected with HSV-1(F) BAC or UL13_K176A_ strains at an MOI of 5 for 16 h. The cells were fixed to perform a LMNA/C immunofluorescence assay. LaminA/C staining was used to define the nuclear boundaries in the cell. FIJI ImageJ software was used to measure nuclear cross-sectional area and perimeter. The nuclear contour ratio was calculated using the formula (4π*area)/(perimeter^2^), as previously described (73). In each sample, fifty nuclei were used to measure the nuclear contour ratio for each experiment.

### Nuclear membrane structural analysis

Images of the nucleus were taken with a Leica DFC7000T confocal microscope using Leica software. Using ImageJ, the freehand tool was used to outline 50 nucleus of Vero cells uninfected or infected with WT, US3_K220A_, and UL13_K176A_. The area was measured in cm^2^ before running the statistical analysis.

### Statistical Analyses

All statistical comparisons were between more than two samples and so were analyzed using one-way analysis of variance (ANOVA) using the Tukey method for multiple comparisons as implemented in GraphPad Prism 8.

## RESULTS

### pUL13 catalytic activity is important to control pUS3’s regulation of NEC and redistribution of Lamin A/C

Mutation of pUS3 in HSV-1 infected cells results in the formation of NEC aggregates and exacerbated disruption of the nuclear lamina at the locations of those aggregations, failure to cause nuclear envelope convolution, and failure of de-envelopment (15, 16). To test if mutation of pUL13 would generate the same phenotypes, we compared the phenotypes of the US3 mutant virus (US3_K220A_) with those of two different isolates of UL13 mutant viruses in which the kinase invariant lysine at residue 176 was changed to alanine (called UL13_K176A_) (Figure1) and confirmed via sequencing. Mutation of pUL13 at K176 has been previously shown to ablate the kinase activity of pUL13 (74). In immunofluorescence assay of Vero cells infected with this virus (Figure 2), the NEC mislocalized in an aggregated pattern at the nuclear rim (Figure 2, C and K), consistent with previous reports (54). However, the NEC aggregation phenotype seen with pUL13_K176A_ differed compared to pUS3_K220A_ in several ways. NEC aggregates in UL13_K176A_ are substantially larger, connected, and reduced in frequency (less prevalent) at the nuclear rim compared to pUS3_K220A_ (Figure 2A, H, and N). In addition, we observed exacerbated nuclear lamina mislocalization in pUL13_K176A_ (Figure 2, D and L) similar to the typical pUS3_K220A_ phenotype (Figure 2, B and I), although the typical perforations observed in the lamina network of pUS3_K220A_ infected cells were larger compared to those seen with pUL13_K176A_ (Figure 2B and I, compared to D and L). Thus, pUL13 catalytic activity is important in the regulation of NEC localization and laminA/C disruption.

**Figure 1.**
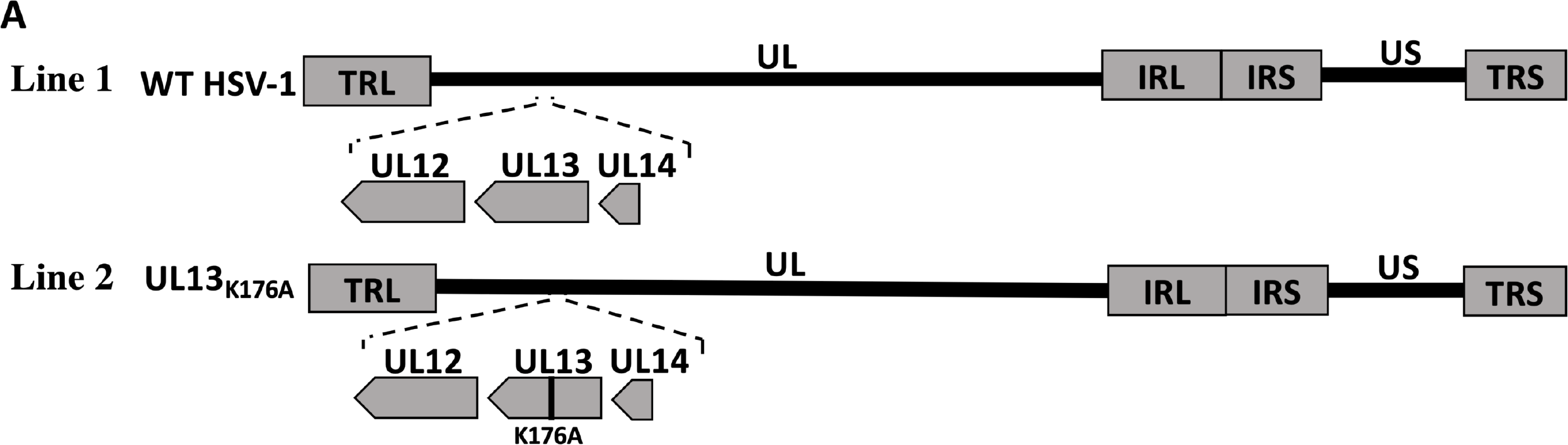
Schematic diagrams of recombinant viruses (A) Schematic diagram of the WT HSV-1(F) BAC-derived genome (line 1) and the UL13_K176A_ recombinant virus constructs (line 2). Line 1 also shows the expanded region of the UL13 locus with its neighboring genes (UL12 and UL14) with respect to the unique long (UL) and unique short (US) sequences that are flanked by the inverted repeats (TRL and IRL, and IRS and TRS) in wild-type. Line 2 shows the UL13_K176A_ virus that has a single amino acid substitution of the invariant lysine (K) to alanine (A) in the ATP-binding domain sequence coding of the UL13 gene.

**Figure 2.**
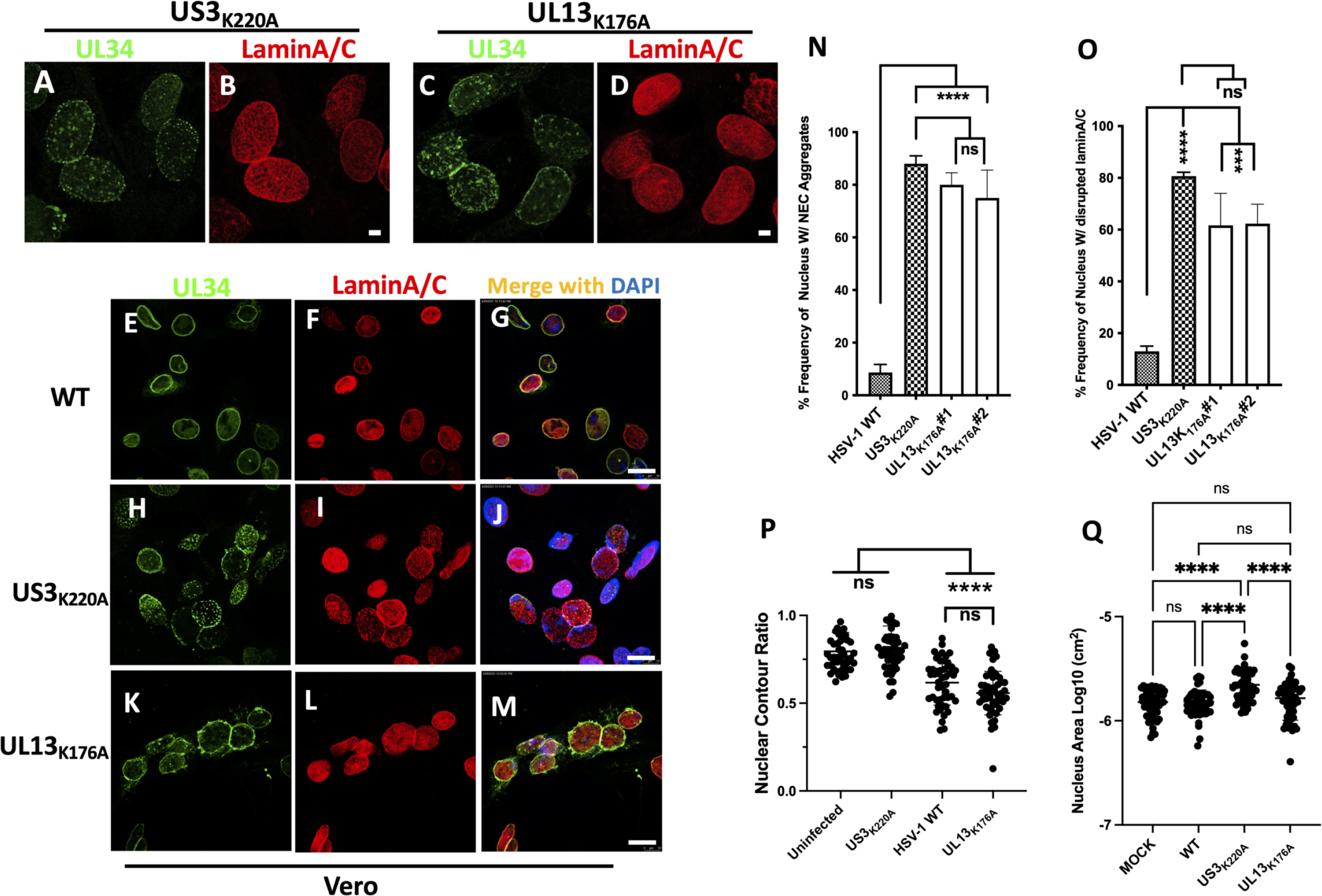
pUL13 catalytic activity is important to regulate NEC aggregation and Lamin A/C disruption (A-I) Representative digital confocal images of infected cells. Vero cells were infected with the WT, US3_K220A_, or UL13_K176A_. Cells were fixed at 16 hpi and stained using antibodies directed against Lamin A/C (green) and pUL34 (red). (A-D) Scale bar = 3μm and (E-M) Scale bar = 10μm. Representative images of one of three independent experiments are shown. (N) and (O) show the frequencies of infected cell nuclei that show NEC aggregates and large gaps in the lamina, respectively. The error bars in each graph represent the range of the values. (P) and (Q) show nuclear contour ratio and nuclear area measurements, respectively, in which each dot indicates the value for an individual infected cell nucleus. The horizontal line indicates the mean value. Statistical significance was determined by one-way analysis of variance (ANOVA) using the Tukey method for multiple comparisons. ns, not significant; *, P ≤ 0.05; ***, P ≤ 0.001; ****, P ≤ 0.001. Data on each graph shows the results for one of at least three independent experiments.

Immunoblot analysis showed an equal amount of pUL34 expression in both HSV-1 WT and pUL13 _K176A_ infected cells (data not shown), suggesting the pUL13 kinase activity does not affect the expression of pUL34, unlike pUL13 homologs in beta and gammaherpesviruses (57). Infection with pUL13 _K176A_ mutant viruses also did not result in the formation of larger, more evenly ovoid nuclei as seen with pUS3 mutants as the majority of pUL13_K176A_ mutants infected cells demonstrated convoluted raisin-resembled nuclear membrane similar to WT (Figure 2P). This suggests pUL13 kinase activity does not play a role in remodeling of the nuclear membrane, unlike pUS3 as pUS3_K220A_ infected cell nuclei show more ovoid shape with enlarged size compared to WT and pUL13 _K176A_ (Figure 2Q).

### TEM analyses of pUL13_K176A_ infected cells

pUS3 mutants are defective in de-envelopment as shown by the presence of intrusions at the INM containing accumulations of primary enveloped virions (PEVs) into the nuclei of infected cells (16). Surprisingly, TEM analysis of UL13_K176A_-infected Vero cells showed no evidence for accumulations of PEVs (Figures 3C and D). The morphology of pUL13_K176A_ nuclei (Figures 3C) was similar to that of WT-infected Vero cells (Figures 3A) with the convoluted nuclear envelope, consistent with our IF assay. Thus, the NEC puncta observed in immunofluorescence images of UL13_K176A_ infected cells do not correspond to accumulations of PEVs. Thus, although there are superficial similarities between the phenotypes of pUL13 and pUS3 mutants, the effects on the nuclear envelope of mutations of these two different kinases are quite distinct.

**Figure 3.**
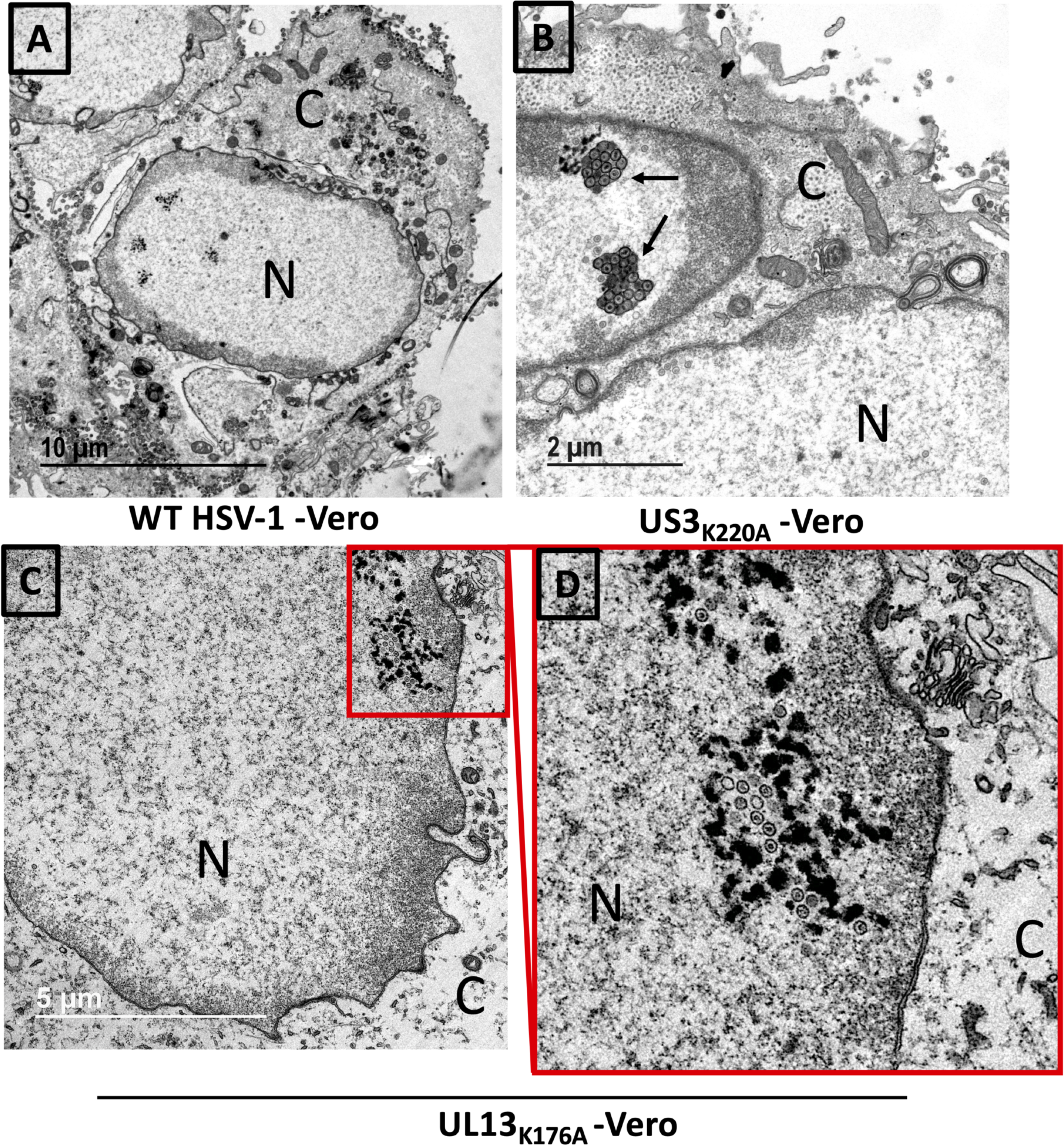
TEM analyses of pUL13_K176A_ infected cells Digital micrographs show Vero cells infected with the US3_K220A_ or pUL13_K176A_ at an MOI of 5 for 20h. Infected Vero cells were fixed with glutaraldehyde, and then processed for TEM. (A) Typical morphology of a WT infected Vero cell. (B) Typical morphology of US3_K220A_ infection with vesicles of enveloped virions. (C-D) morphology of UL13_K176A_ which is similar to the typical morphology of WT infected Vero cells with encapsidated virions in the nucleus, in the cytoplasm, or on the cell surface. Magnification in D 5,400X.

### The phosphorylation status of selected US3 substrates in UL13_K176A_ infected cells

The report from Kato et al. demonstrating that a UL13 deletion affected one function of pUS3 (NEC localization) but not another (inhibition of apoptosis), suggested that phosphorylation of pUS3 might affect its ability to phosphorylate some subset of its substrates. Considering that the NEC is a nuclear membrane-associated physiological substrate of pUS3, while pro-apoptotic proteins are the soluble cytoplasmic physiological substrate of pUS3, we hypothesized that UL13 may regulate the ability of pUS3 to phosphorylate specific classes of its substrates.

To test this hypothesis, chose known pUS3 substrates in three different classes and tested their phosphorylation in UL13_K176A_ infected cells (Figure 4). These included a nuclear membrane- associated proteins (emerin) (26), a cytoplasmic soluble protein (PKA regulatory type II alpha- subunit (PRKAR2A)) (36); and a cytoplasmic membrane-associated protein, (glycoprotein B (gB)) (50). The phosphorylation of emerin and PRKAR2A by pUS3 has been shown to cause a mobility shift in SDS-PAGE (26), so we determined whether the mobility of these proteins was affected in cells infected with wild-type and US3 and UL13 mutant viruses (Figure 4). In Figure 4A, hyperphosphorylation of emerin by pUS3 was shown by the presence of lower mobility bands in WT virus-infected samples that disappeared in US3_K220A_, consistent with previous reports. Hyperphosphorylation of emerin in UL13_K176A_ infected cells was indistinguishable from WT, suggesting that pUL13-mediated phosphorylation of pUS3 is not required for phosphorylation of all pUS3 nuclear substrates. Similarly, consistent with the previous report, we observed a slower migrating form of PRKAR2A in the WT-infected sample that was absent in US3_K220A_-infected cells (Figure 4D). As with emerin, the WT and UL13 mutant migration patterns for PRKARA2A were indistinguishable. pUS3- mediated phosphorylation of gB on its cytoplasmic tail at T887 does not result in an observable mobility shift, but this phosphorylation site is very similar to sites that are phosphorylated by protein kinase A (PKA) and can be detected with an antibody that reacts with phosphorylated PKA substrates (50). gB was immunoprecipitated from cells infected with UL13_K176A_ and total phosphorylated gB was detected using antibodies to gB and to phosphorylated PKA substrates. As observed previously, phosphorylated gB can be detected in WT-infected cells, but is greatly diminished in US3_K220A_ (Figure 4B and C). As with emerin and PRKAR2A the amount of phosphorylated gB in UL13- mutant infected cells is the same as WT. The pUS3 kinase motif (RnX(S/T)), where n ≥ 2 overlaps with some cellular protein kinase phosphorylation motifs, specifically PKA (36) and protein kinase B (Akt) (49), and pUS3, PKA, and Akt can recognize and phosphorylate some of the same substrates (36, 75). The pUL13 consensus phosphorylation motif has been characterized as (A/GSPA/G), where the SP in the center of the motif is invariant (76, 77). This motif does not match the PKA or Akt kinase motif, and it is unlikely that UL13 phosphorylates PKA or Akt substrates. To examine the phosphorylation of pUS3 substrates whose phosphorylation sites match those of PKA and Akt; whole cell lysates were prepared from either mock-infected or cells infected with WT, US3_K220A_, or UL13_K176A_, blotted, and probed with specific phospho-PKA or phospho-Akt substrates antibody (Figure 4E). The pattern of reactive proteins is quite different in WT and US3_K220A_-infected cells, with prominent phosphorylated species observed in WT that are absent in US3_K220A_. However, the phosphorylation patterns of PKA or Akt substrates in pUL13_K176A_ were very similar to WT, suggesting that most pUS3 substrates are phosphorylated in a pUL13-independent manner.

**Figure 4.**
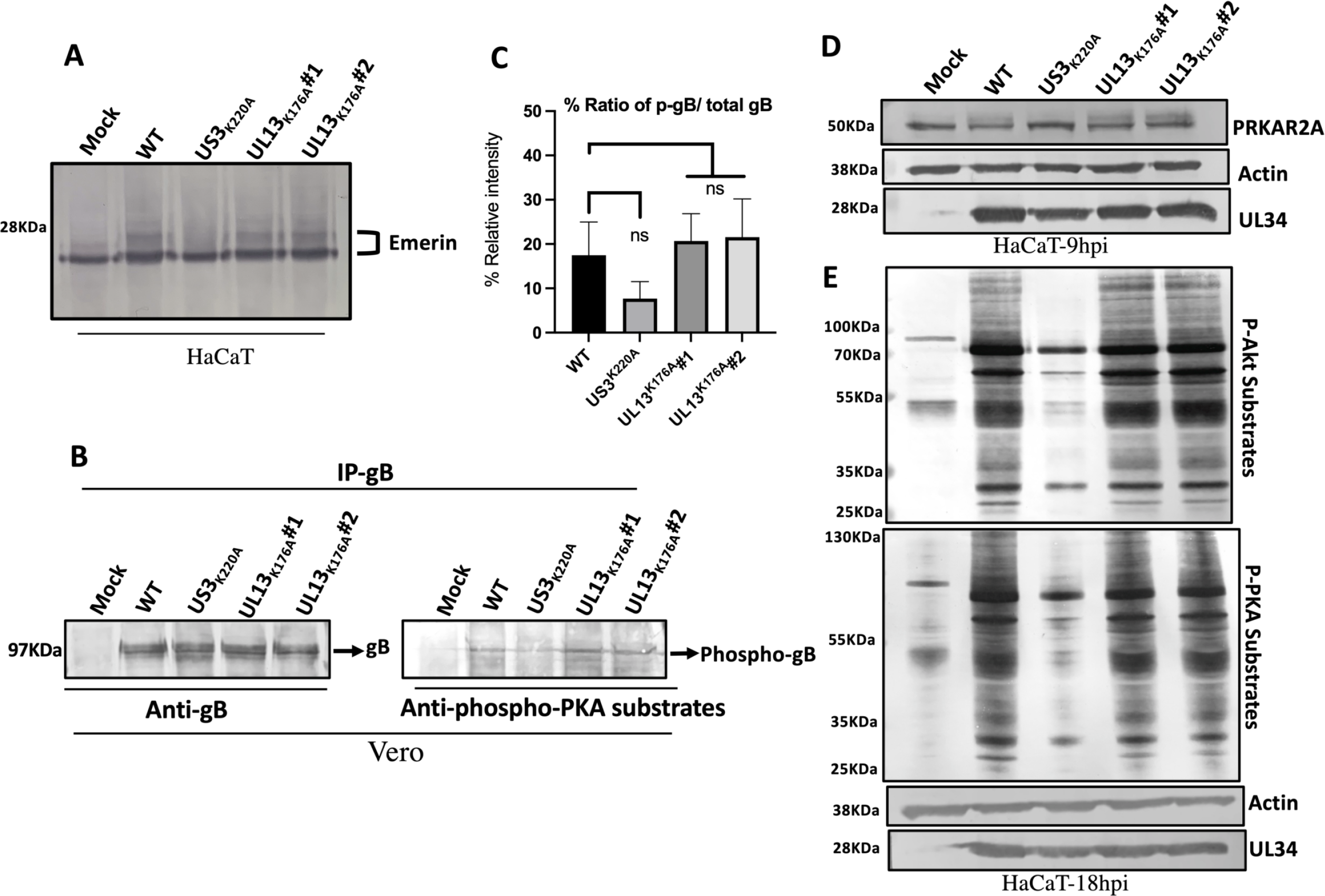
The phosphorylation status of selected US3 substrates in UL13K176A infected cells (A) Emerin hyperphosphorylation assay. Digital images of an immunoblot of nuclear membrane proteins from HaCaT cells infected for 16 h with 5 PFU/cell of BAC-derived WT HSV-1(F), US3_K220A_ or pUL13_K176A_ or uninfected and probed for emerin. (B) gB immunoprecipitation assay. Vero cells were uninfected or infected with 5 PFU/cell of BAC-derived WT HSV-1(F), US3_K220A_ or pUL13_K176A_. After 18 h of infection, cell lysate samples were immunoprecipitated with the anti- gB antibody. Immunoprecipitated protein was separated by SDS-PAGE, blotted onto nitrocellulose membrane, and probed with either anti-gB antibody (left panel) to detect total gB population or anti-phospho-PKA substrate antibody (right panel) to detect US3-dependent phosphor-gB. (C) gB densitometry. The relative density of phosphorylated gB proteins was normalized to that of total gB proteins shown in (B). ns, not significant. (D) Digital images of an immunoblot of protein lysates from HaCaT cells infected for 9 h with 5 PFU/cell of BAC-derived WT HSV-1(F), US3_K220A_ or pUL13_K176A_ or uninfected and probed for PRKAR2A (top panel), actin (middle panel; equal loading control) or pUL34 (bottom panel; equal infection control). (E) Digital images of immunoblots for phosphorylated Akt (top panel) and PKA (second-from-top) motifs in lysates of HaCaT cells infected for 18 h with 5 PFU/cell of BAC-derived WT HSV-1(F), US3_K220A_ or pUL13_K176A_. Probes for actin (third from top) and pUL34 (bottom) served as controls for equivalent loading and infection, respectively.

### UL13 phosphorylation motifs in pUS3 are not important for regulating NEC localization

Kato et al., reported a correlation between phosphorylation of pUS3 by pUL13 and regulation of NEC localization but did not show that these events were causally connected. To address this question, we identified matches in the pUS3 sequence to the reported pUL13 consensus phosphorylation motif (SP) and changed the phosphorylatable residues within all of these motifs to alanine. Analysis of the pUS3 sequence along with Alpha-fold modeling revealed four such motifs (Table 1 and Figure 5). Furthermore, one of these (S139) has been shown to be phosphorylated in HSV-infected cells (77). To test the significance of the four putative pUL13 phosphorylation sites, we created recombinant viruses in which each of the serine residues was substituted with alanine and one in which all four were substituted at the same time. Two independent isolates of each virus were tested to ensure that any phenotype was due to the intended mutation (Figure 6). Cells infected with these viruses were tested for altered NEC localization (Figure 7A-H) and for exacerbated lamina disruption (data not shown). None of the putative pUL13 phospho-site mutations reproduced the phenotype of the pUL13_K176A_ mutation consistent with the quantification (Figure 7I), suggesting that the pUL13 effect on NEC distribution is not due to phosphorylation of pUS3. Kato et al., reported that deletion of pUL13 resulted in a mobility shift in pUS3, consistent with loss of phosphorylation (54). We tested for mobility shift of pUS3 in cells infected with pUL13_K176A_ and with the pUL13 putative phospho-site mutants (Figure 7J). We observed no difference in migration of pUS3 in any of these conditions, suggesting that pUL13 catalytic activity does not mediate the phosphorylation of any large fraction of pUS3 molecules. Noteworthy, all the pUS3 mutants were expressed to the same degree as the native pUS3 (Figure 7J).

**Figure 5.**
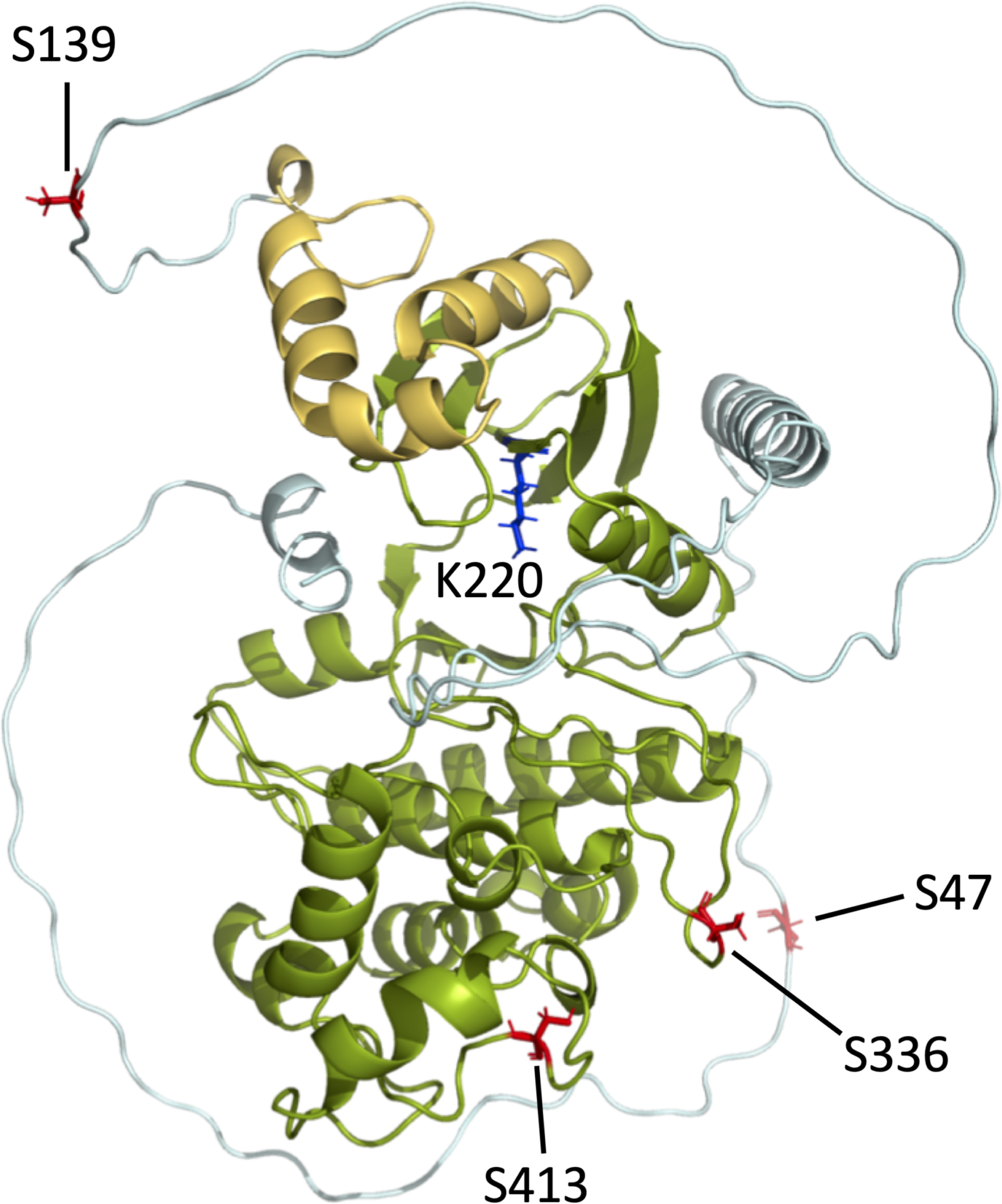
Alpha-fold modeling of the pUS3 structure Shown is a typically folded protein kinase domain at the C-terminus (green), an N-terminal region consisting of residues 1-191 that is largely disordered (sky blue), with a small ordered domain (gold) just preceding the kinase domain. The S47, S139, S336, and S413 (red) represent sites of serine residues that match the pUL13 consensus phosphorylation motif (SP). K220 (blue) is the invariant lysine residue in the ATP-binding site domain. The S336 residue is predicted to be within the putative protein kinase domain. S47, S139, and S413 are found in predicted disordered regions of pUS3. All four of the SP motifs are predicted to be accessible.

**Figure 6.**
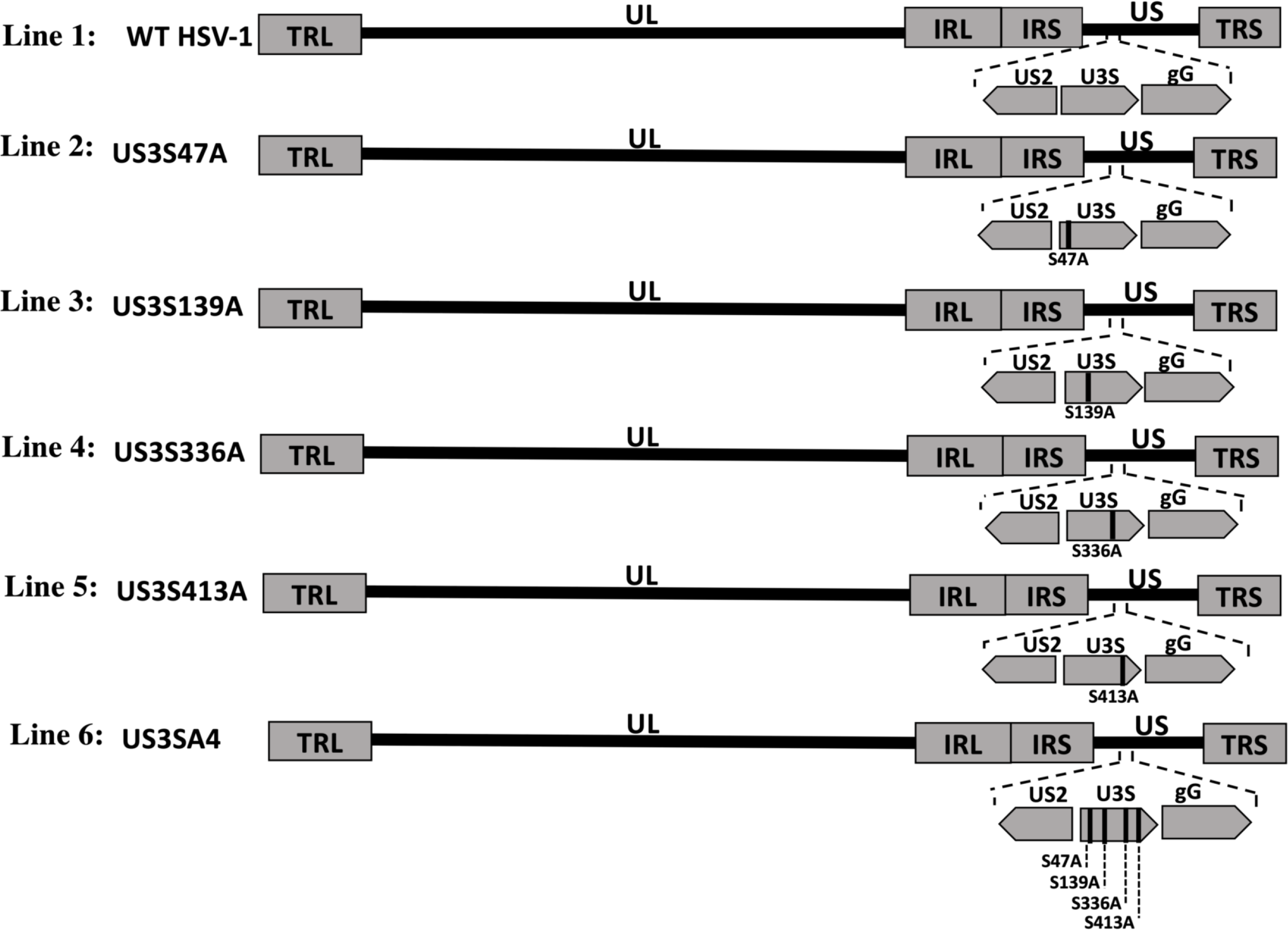
Schematic diagrams of US3 recombinant viruses Schematic diagram of the WT HSV-1(F) BAC-derived genome (line 1), and the US3 (line 2-6), recombinant virus constructs. Line 1 shows the expanded region of the US3 locus with its neighboring genes (US2 and US4) with respect to the unique long (UL) and unique short (US) sequences that are flanked by the inverted repeats (e.g. TRL and IRL, and IRS and TRS) in wild-type. Line 2-5 show the US3 virus that has a single amino acid substitution of the serine (S) to alanine (A) in the SP domain sequence coding of the US3 gene designated as US3_S47A_, US3_S139_, US3_S336A_ and US3_S413A_, respectively. Line 6 shows the US3 virus that has all amino acid substitution of the serine (S) to alanine (A) in the SP domain sequences coding of the US3 gene designated as US3_SA4_.

**Figure 7.**
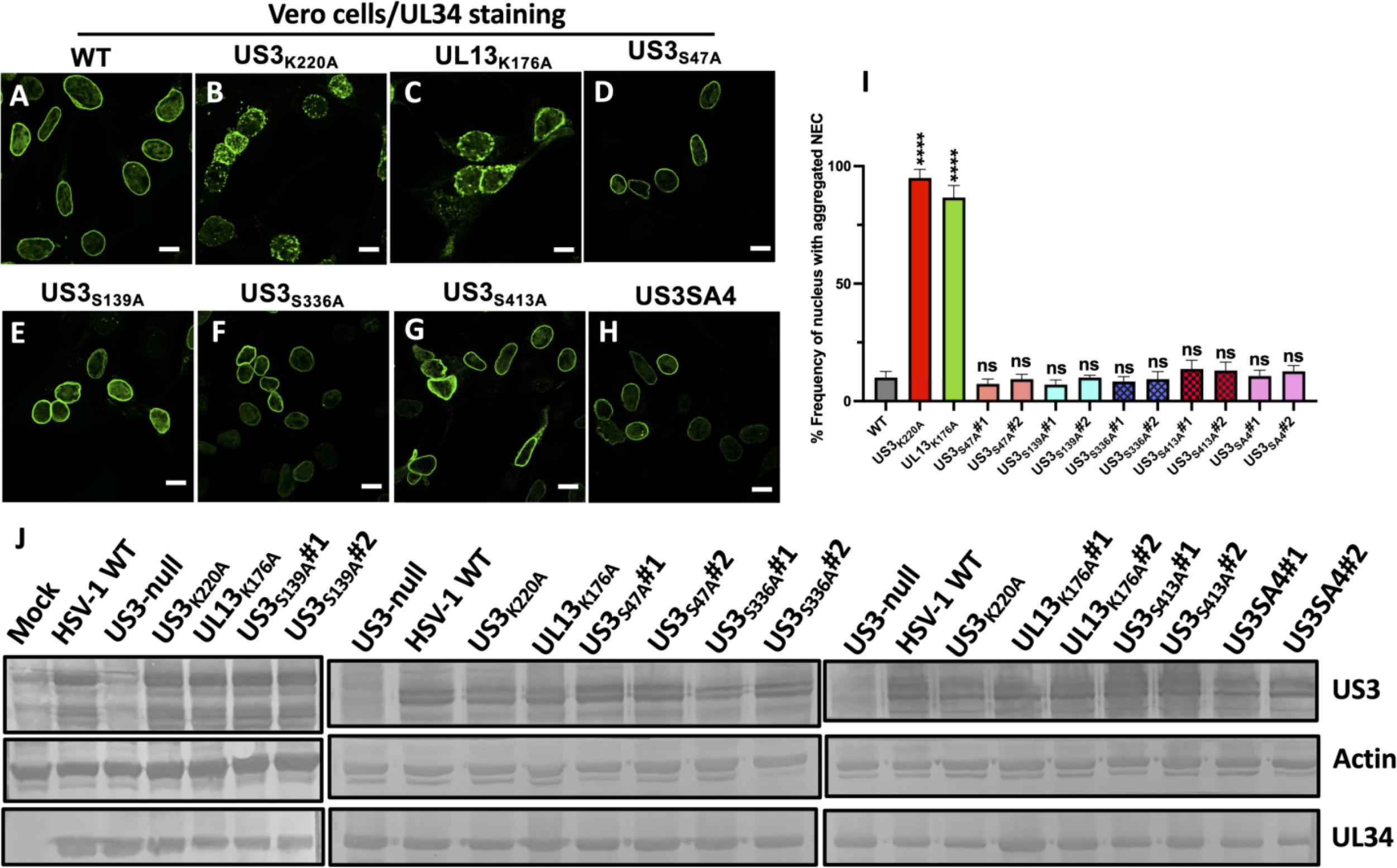
Nuclear egress phenotypes comparison of pUL13_K176A_ and pUS3 mutants (A-H) Shown are digital confocal images representing the localization of UL34 in infected Vero cells Green: pUL34. Cells were infected with HSV-1(F) (BAC), US3_K220A_, UL13_K176A_, and US3 mutants for 16 h at a MOI of 5, fixed at 16 hpi, and stained using antibodies directed against pUL34. Scale bar = 10μm. Representative images of one of three independent experiments are shown. (I) Frequencies of infected cell nuclei that show NEC aggregates. The error bars show the range of the values. Statistical significance was determined by one-way analysis of variance (ANOVA) using the Tukey method for multiple comparisons. ns, not significant; ***, P ≤ 0.001. Data on graph represents at least three independent experiments. (J) US3 mobility shift assay of UL13_K176A_ and US3 mutants. Digital images of immunoblots for the indicated proteins in lysates of Vero cells infected for 18 h with 5 PFU/cell of BAC-derived WT HSV-1(F), US3_K220A_ or pUL13_K176A_ or US3 mutants. UL34 serves as a marker of equivalent infection and actin as a loading control.

**Table 1.**
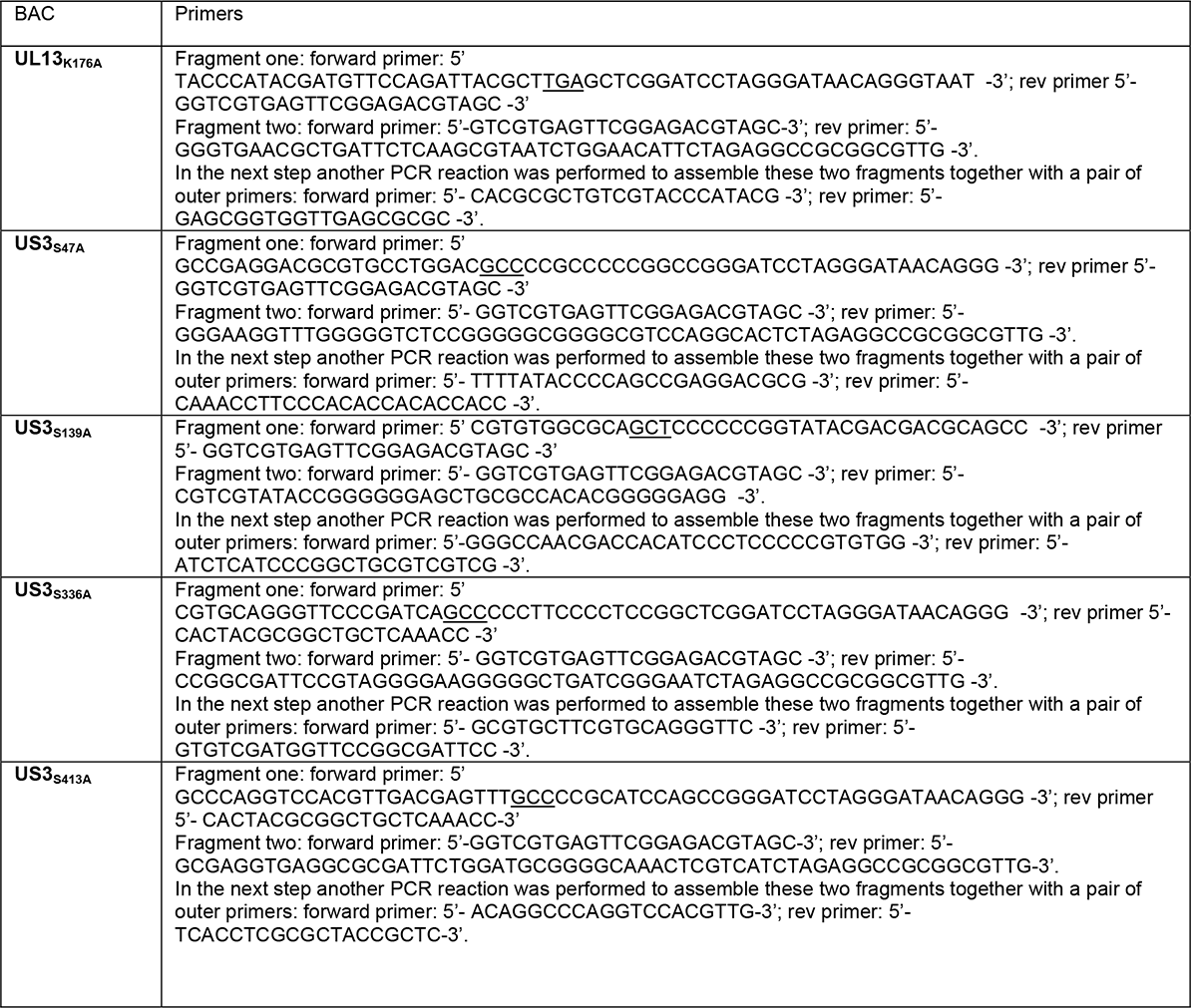
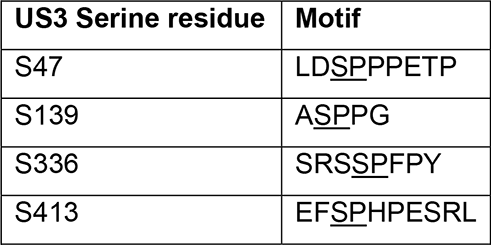
UL13 phosphorylation motif sites on US3

### Growth analysis of pUL13K176A and pUS3 S to A mutants on HaCaT and SH- SY5Y

pUS3 activity is not essential for viral growth in cell culture and the magnitude of the growth defect associated with US3 mutations is cell type-dependent. We have observed a large growth defect associated with loss of pUS3 catalytic activity in the HaCaT keratinocyte and SH- SY5Y neuronal cell lines, which represent cell types important to HSV pathogenesis. The significance of UL13 mutation in these cell types has not previously been determined. To test the significance of pUL13 kinase function and of the putative pUL13 phosphorylation sites of pUS3 in viral growth, HaCaT and SH-SY5Y cells were infected at an MOI 5 and at 16 hpi, and the viral infectivity in the culture was measured by plaque assay on Vero cells. SSG analyses in both HaCaT (Figure 8) and SH-SY5Y (data not shown) cells demonstrated a growth defect phenotype for all the US3 serine to alanine single amino acid substitution and US3SA4 mutants, with both cell lines showing ∼100-fold reduced growth of the US3_S47A_ and US3SA4 mutant compared to the wild-type and UL13_K176A_ (Figure 8) and ∼10 to 50-fold reduced growth of the US3_S139A_, US3_S336A_, and US3_S413A_ mutants. In contrast, the UL13_K176A_ mutant showed unimpaired growth in both cell lines, suggesting that the growth defects associated with the US3 point mutants resulted from intrinsic effects on pUS3 function rather than effects due to pUL13 mediated phosphorylation.

**Figure 8.**
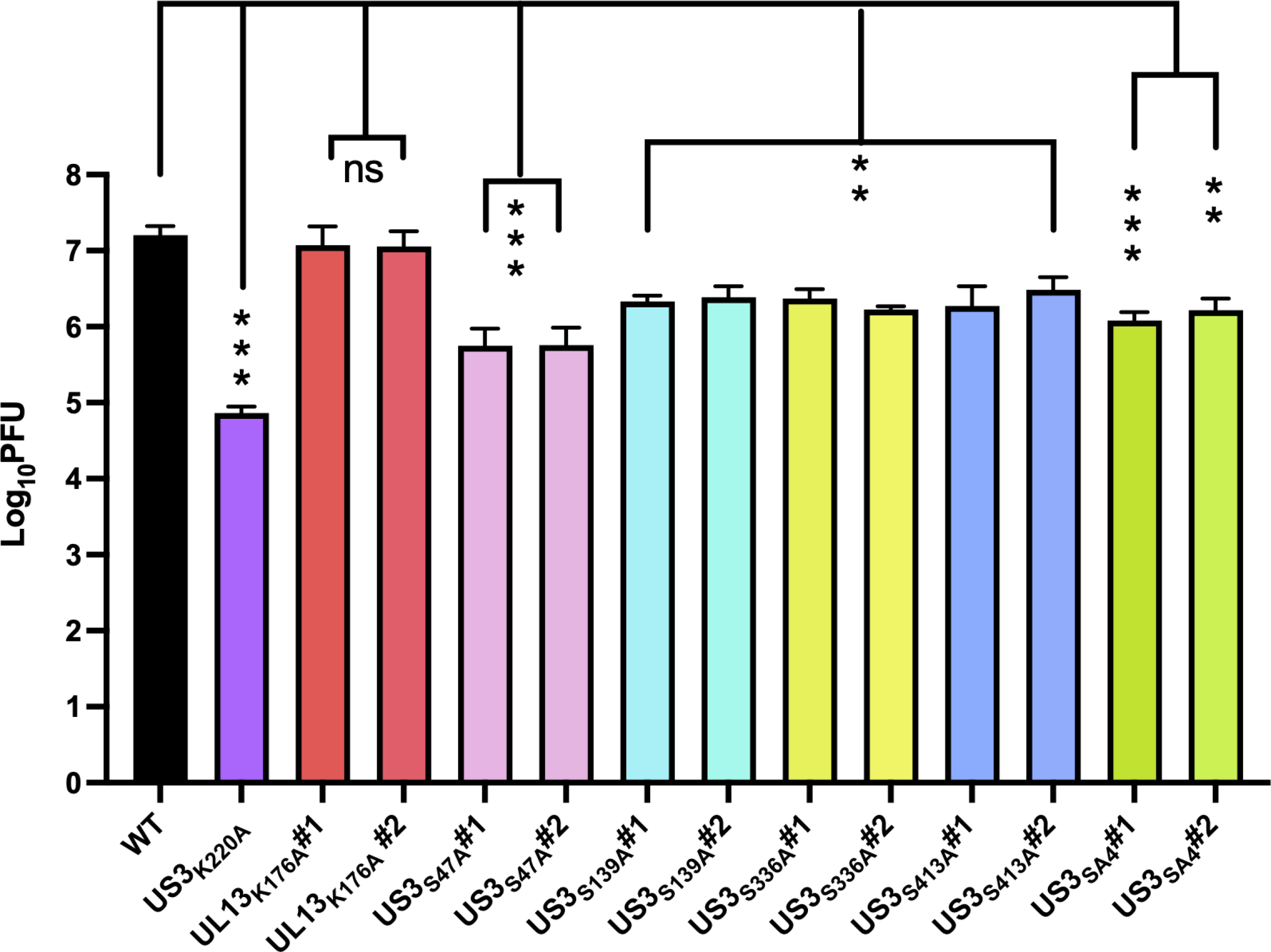
SSG analysis of US3 mutants on HaCaT Shown are single-step growth analyses of HaCaT cells that were infected with WT, US3_K220A_, UL13_K176A_, US3_S47A_, US3_S139A_, US3_S336A_, US3_S413A_, or US3SA4 viruses at an MOI 5. At 16 hpi cells were frozen, thawed, and titered on Vero cells. One of two independently performed experiments consisting of three biological replicates is shown. The error bars in each graph are represented as mean with range. Statistical significance was determined by one-way analysis of variance (ANOVA) using the Tukey method for multiple comparisons performed on log- transformed data. ns, not significant; *, P ≤ 0.05; ***, P ≤ 0.001.

### UL13 colocalizes with pUL31 inside the nucleus and interacts with pUL31

Co- expression of pUL31 and pUL34 in the absence of other viral proteins results in the formation of NEC aggregates at the nuclear rim (6). We previously reported that US3 is sufficient to smooth out the UL31/UL34 self-assembled organizations at the nuclear rim in a co-transfected assay (33). To test if HSV-1 UL13 is similarly sufficient to prevent the NEC self-association, we determined the localization of epitope-tagged NEC components and of EGFP-pUL13 in Vero cells co-transfected with EGFP-UL13, FLAG-UL31, and HA-UL34 plasmids constructs and fixed 48 h post-transfection (Figure 9). As previously reported, co-expression of pUL31 and pUL34 in the absence of UL13 resulted in co-localization of pUL34 and pUL31 and in the formation of NEC self-assembled organizations on the nuclear rim (Figure 9E and F). pUL13 nuclear localization was also observed (Figure 9A) in the absence of other viral proteins consistent with previous reports for pUL13 homologs (78, 79). Interestingly, the expression of pUL13 along with pUL34 and pUL31 resulted in some even distribution of the NEC at the nuclear membrane (Figure 9J), similar to that seen in the cell transiently expressing pUS3 along with pUL34 and pUL31. Surprisingly, however, a portion of pUL31, but not pUL34 also colocalized with UL13 inside the nucleus in large aggregations, suggesting a physical interaction between pUL31 and pUL13 (Figure 9I). Interestingly, although pUL31 was partially recruited away from the nuclear rim when pUL13 was co-expressed, pUL34 remained confined to the nuclear rim, suggesting that there was some excess pUL31 in the nucleus. The interaction between pUL31 and pUL13 in transfected cells was confirmed by co-immunoprecipitation of pUL13 with pUL31-FLAG (Figure 9L), suggesting that pUL31 is a novel interaction partner for pUL13.

**Figure 9.**
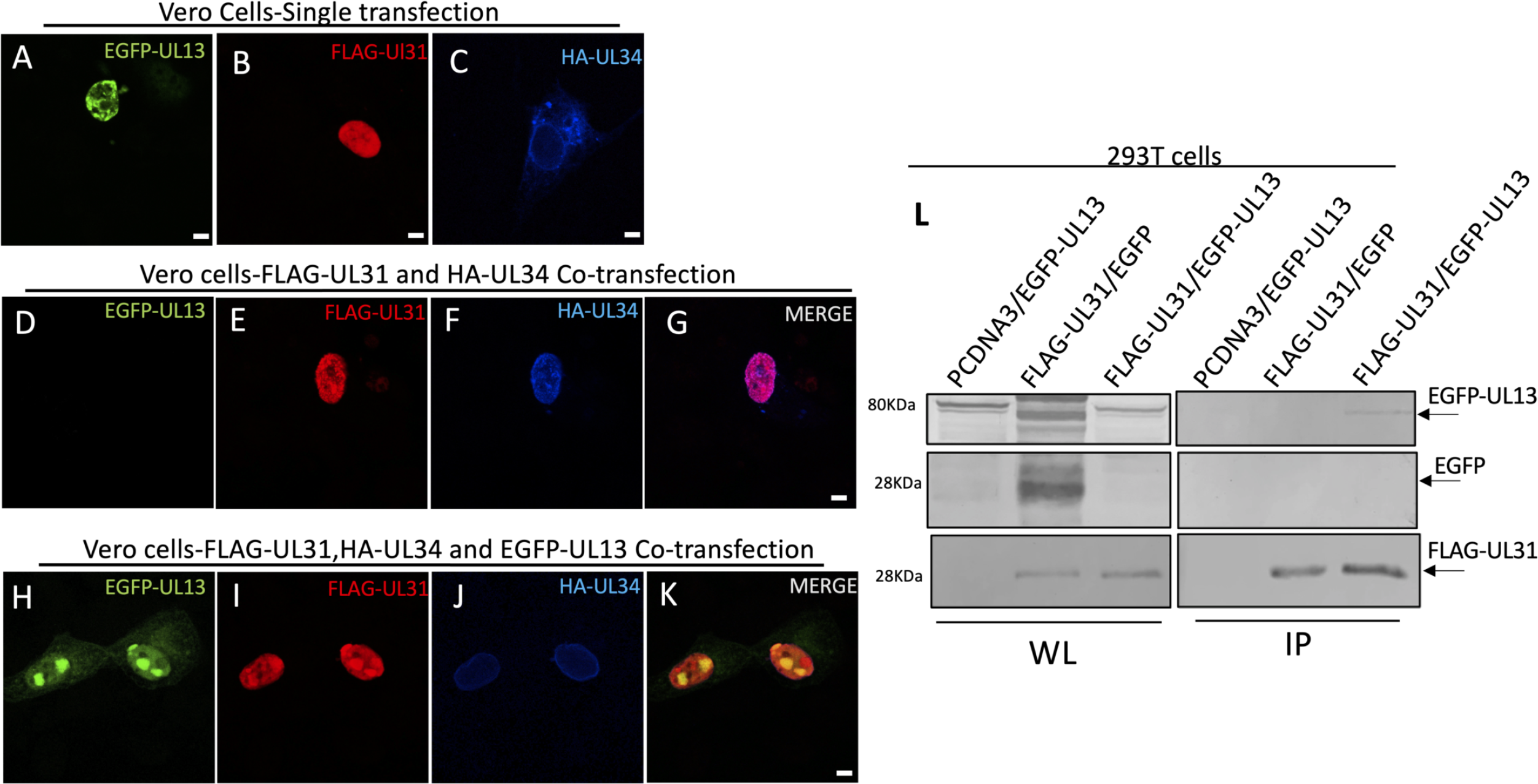
UL13 colocalizes with pUL31 inside the nucleus and interacts with pUL31 Shown are digital confocal images of representing the localization of FLAG-UL31, HA-UL34, and EGFP-UL13 transiently co-expressed proteins in Vero cells, single-cell transfection (A-C), without EGFP-UL13(D-G), or with EGFP-UL13 co-expression (H-K). Cells were fixed at 48 h post-transfection and were stained using antibodies directed against HA and FLAG. Green: EGFP-UL13, Blue: HA-UL34, Red: UL31-FLAG. Representative images of one of three independent experiments are shown. Bars, 5μm. (L) Co-immunoprecipitation of EGFP-UL13 and FLAG-UL31. 293-T cells were co-transfected with FLAG-UL31 and EGFP-UL13 plasmids. Cells were lysed after 48 hours post-transfection. Cell lysate samples were collected and incubated with FLAG magnetic beads to immunoprecipitate UL31 overnight. The whole lysate (WL) samples and immunoprecipitated (IP) eluents were separated by SDS-PAGE, blotted onto nitrocellulose membrane, and probed as indicated in the figure.

## DISCUSSION

The tight curvature of biological membranes is energetically unfavorable and must be coupled to energetically favorable events in order to occur. Spontaneous budding of artificial membranes induced by exposure to purified NEC complexes suggests that spontaneous self- association of the NEC to a higher-order structure is an energetically favorable event that can drive nuclear membrane budding during herpesvirus nuclear egress (12, 13, 33). The formation of hexameric arrays in NEC crystal structures and in vitro budded vesicles, as well as the formation of ordered NEC organization in PEVs all support a central role for NEC self- association as a critical event in membrane budding (14). In wild-type virus-infected cells, however, membrane budding is a rare event in the absence of capsid, suggesting that the NEC self-association is negatively regulated in the process of membrane budding. Several lines of evidence have suggested that pUS3 phosphorylation of pUL31 is one regulator of that self- association. In cells infected with pUS3 mutant viruses, the NEC forms distinctive punctate aggregates in the nuclear envelope, and in US3 mutant infected cells, these aggregates are associated with inclusions of the inner nuclear membrane that contain multiple PEVs (16). Similar structures are observed when US3 phosphorylation motifs in pUL31 are mutated (19). Furthermore, we previously reported that pUS3 controls the formation of NEC aggregates even when capsid envelopment is prevented by failure to express the major capsid protein and, in this circumstance, NEC aggregation is associated with the formation of folded and tightly curved regions of the nuclear envelope (33). Regulation of NEC self-association for budding by pUS3- mediated phosphorylation is also suggested by the apparently opposing effects of pUS3 and a phosphatase recruited to the NE by pUL21 (80). Thus, loss of pUS3-mediated regulation of NEC self-association for membrane curvature might contribute to both NE deformation in US3/capsid protein double mutants, and the impairment of de-envelopment in US3 single- mutants. Inhibition of NEC self-association by phosphorylation might prevent capsid-less, unproductive budding and phosphorylation of the NEC might promote capsid release in de- envelopment.

A previous study by Kato et al., suggested that pUL13 might provide an additional level of regulation, controlling pUS3 NEC phosphorylation function by phosphorylation of pUS3 (54). The results reported here, however, suggest a different role for pUL13 in the regulation of the NEC. Catalytically inactivating mutations of both pUS3 and pUL13 induce formation of NEC aggregates in the nuclear envelope, and in both cases, these aggregates apparently induce clearance of the lamin network, possibly by recruitment of other factors, including cellular protein kinases, that disrupt lamina structure (81, 82). Nonetheless, the aggregates formed in pUL13- and pUS3-mutant infections are different both in appearance and functional consequence. Areas of NEC association in pUS3-mutant infected cells are both smaller, more discrete, and more numerous than those formed in pUL13 mutant infection. Most notably, mutation of pUL13 does not result in accumulation of PEVs suggesting that it does not regulate de-envelopment, nor is pUL13 mutation associated with the loss of nuclear envelope convolution and expansion of nuclear volume also associated with pUS3 mutations. These results suggest that these two viral protein kinases are involved in two different pathways of controlling the NEC. Our results are consistent with the hypothesis that there are at least two modes of NEC self-association. One of these modes is membrane curvature-inducing and is regulated by pUS3 phosphorylation of the pUL31 N-terminus. The other mode of self- association does not induce membrane deformation and is regulated by pUL13. The significance of this second form of self-association is unclear. It is possible that it represents an unproductive aggregation of the NEC and that pUL13 serves to minimize the diversion of NEC heterodimers into these unproductive complexes. This suggestion could be tested by examining the effects of pUL13 and pUS3 in *in vitro* NEC budding assays (14), where added pUS3 might be expected to inhibit budding and pUL13 might promote it. Double mutation of pUS3 and pUL34 has shown that some, but not all of the pUS3 effect on virus replication in HaCaT cells is due to its regulation of nuclear egress (33). The lack of a growth defect for the pUL13 mutant tested here shows that, at least in HaCaT cells, pUL13-dependent regulation of NEC localization and association is not necessary for nuclear egress and does not significantly promote virus production. Despite the lack of the growth defect for UL13_K176A_ mutants, it reproduced the UL13-null phenotype with respect to NEC distribution, confirming that mutation of K176 to A176 in UL13 did indeed inactivate the catalytic activity of the pUL13.

It was surprising that mutation of pUL13 had no effect on phosphorylation of any of the substrates specifically tested here or on the range of pUS3 substrates that overlap with PKA and Akt substrates. While Kato et al, showed UL13-dependent phosphorylation of pUL31, they did not show that this phosphorylation was mediated by pUS3. As a result, there is currently no evidence that pUL13 controls pUS3-mediated phosphorylation of any substrate. The pUL31 sequence contains three SP motifs that might be phosphorylated by pUL13, including two near the N-terminus at residue S11 and S43, suggesting the possibility that pUL31 might be the relevant substrate of pUL13 for regulation of NEC localization/aggregation.

Our results are also inconsistent with those of Kato et al., in that we observed no shift in pUS3 migration in SDS-PAGE following mutation of pUL13 or of possible pUL13 phosphorylation sites in pUS3. It is possible that differences in experimental conditions masked this effect in our hands, but our observations suggest that, if pUL13 phosphorylates pUS3, this does not affect pUS3 function in nuclear egress and might regulate one or more of the many other functions of pUS3. Interestingly, mutation of each of the pUS3 SP motifs significantly impaired virus replication in HaCaT cells, even though mutation of pUL13 did not. This suggests that these residues participate in UL13-independent functions of pUS3. The tertiary structure of pUS3 is undetermined, but only one of these residues (S336) is within the putative protein kinase domain. Alpha-fold modeling of the pUS3 structure (Figure 5) shows a typically folded protein kinase domain at the C-terminus and an N-terminal region consisting of residues 1-191 that is largely disordered with a small ordered domain just preceding the kinase domain. S47, S139, and S413 are found in predicted disordered regions of pUS3 that may control its interactions with other proteins in the infected cell. All four of the SP motifs are predicted to be accessible.

The co-localization of pUL13 and pUL31 in large, spherical intranuclear inclusions in transfected cells is very similar to the pUL31 localization observed in cells in which the DNA damage response has been induced. It is also possible that pUL13 interaction with pUL31 is mechanistically related to the defined role of pUL31 in DNA damage response (80), in which pUL31 antagonizes the pUL13 induction of DNA damage (58, 83). Future studies are warranted for a more mechanistic explanation for this novel interaction in the infection system.

In conclusion, we have provided data showing that pUL13 regulates NEC localization in a manner distinct from, and probably independent of pUS3 and that this regulation does not affect virus de-envelopment. Rather, our results are consistent with a model in which the HSV NEC can associate in at least two ways, each of which is regulated by one of these two kinases. Finally, a physical and possibly functional interaction between pUL13 and pUL31 may affect a host cell response to virus infection.

## Acknowledgments

The authors would like to thank Alison Haugo-Crooks and Shaowen White for critical reading of the manuscript. This work was supported by NIH grant R21AI133155 and by the Department of Microbiology and Immunology.

